# Metabolomic signatures of coral bleaching history

**DOI:** 10.1101/2020.05.10.087072

**Authors:** Ty N.F. Roach, Jenna Dilworth, H. Christian Martin, A. Daniel Jones, Robert Quinn, Crawford Drury

## Abstract

Coral bleaching, a process where corals expel their photosynthetic symbionts, has a profound impact on the health and function of coral reefs. As global ocean temperatures continue to rise, bleaching poses the greatest threat to coral reef ecosystems. Here, untargeted metabolomics was used to analyze the biochemicals in pairs of adjacent corals from a patch reef in Kāne‘ohe Bay, Hawai‘i, where one colony in the pair bleached (in 2015) and recovered while the other did not bleach. There was a strong metabolomic signature of prior bleaching history four years after recovery found in both the host and its algal symbionts. Machine learning analysis determined that the strongest metabolite drivers of the difference in bleaching phenotype were a group of betaine lipids. Those with saturated fatty acids were significantly enriched in thermally tolerant corals and those with longer, unsaturated and diacyl forms were enriched in historically bleached corals. Host immune response molecules, Lyso-PAF and PAF, were also altered by bleaching history and were strongly correlated with symbiont community and algal-derived metabolites suggesting a role of coral immune modulation in symbiont choice and bleaching response. To validate these findings, we tested a separate *in situ* set of corals and were able to predict the bleaching phenotype with 100% accuracy. Furthermore, corals subjected to an experimental temperature stress had strong phenotype-specific responses in all components of the holobiont, which served to further increase the differences between historical bleaching phenotypes. Thus, we show that natural bleaching susceptibility is simultaneously manifested in the biochemistry of the coral animal and the algal symbiont and that this bleaching history results in different physiological responses to temperature stress. This work provides insight into the biochemical mechanisms involved in coral bleaching and presents a valuable new tool for resilience-based reef restoration.

## Introduction

Reef building corals form an obligate endosymbiosis with single-celled photosynthetic microorganisms known as Zooxanthellae. This partnership is one of the oldest known mutualistic symbioses, dating back more than 160 million years (LaJeunesse et al. 2018). Despite the tight coupling in evolutionary fitness between the two organisms, environmental stressors such as increased temperature can cause the breakdown of this symbiosis in a process known as coral bleaching. As zooxanthellae typically provide upwards of 90% of the corals’ energy (Muscatine and Porter 1977), bleaching frequently leads to substantial coral mortality, compromising the structure and function of coral reef ecosystems.

Thermally-induced bleaching is occurring with increased frequency and severity, and is expected to impact most of the world’s reefs annually by mid-century, sparing few temporary refugia (van Hooidonk, Maynard, and Planes 2013). As temperature stress threatens the long-term persistence of reef ecosystems, there is a pressing need to develop an integrative understanding of the holobiont response to thermal stress and the interactions between different components of the symbiosis. This could facilitate the ability to rapidly measure changes in coral physiology and support active intervention strategies (Ocean Sciences Board 2019). Historically, investigating the bleaching response in corals has focused on molecular markers that describe patterns of genes, transcripts, proteins, and metabolites during temperature stress. For example, host gene expression (Barshis et al. 2013; Palumbi et al. 2014; Bay and Palumbi 2015), the bacterial microbiome (Grottoli et al. 2018; Ziegler et al. 2017) and the metabolome (Sogin et al. 2016) have all been shown to change under elevated temperatures. However, our understanding of longer-term impacts of past bleaching is limited (Pinzón et al. 2015; Thomas and Palumbi 2017), and differences between thermally resistant and susceptible phenotypes in unstressed conditions is largely unexplored. Corals exhibit a broad range of thermal tolerance (Howells et al. 2016; Matsuda et al. 2019), which can be used as a framework to investigate the dynamics of resilience and leveraged for restoration. The identification of the drivers and latent effects of bleaching are fundamental for restoration efforts (van Oppen et al. 2017) and development of tools for resilience-based management (Anthony et al. 2015) under a changing climate.

Recently, mass spectrometry-based metabolomics approaches have allowed researchers to explore the chemical diversity of a wide range of complex biological systems (da Silva, Lopes, and Silva 2017). Here, we applied an untargeted metabolomics approach to examine the effects of past bleaching and experimental heat stress on the coral holobiont. We sampled pairs of adjacent corals with different historical bleaching phenotypes on a patch reef in Kāne‘ohe Bay, Hawai‘i four years after a bleaching event (Figure S1) (R Cunning, Ritson-Williams, and Gates 2016; Matsuda et al. 2019) and identified a strong signature of bleaching history in the metabolomic data that was primarily driven by the abundance of a class of highly saturated betaine lipids in resilient corals. Host immune response molecules were also significantly different between bleaching phenotypes and correlated strongly with algal-derived metabolites. After re-exposing these corals to thermal stress, we found phenotype-specific heat responses attributed to the symbiotic algae, the coral host, and putative bacterial natural products, which magnified the prior differences between historical bleaching phenotypes.

## Methods and Materials

### Coral collection, stress test, and metabolomic sampling

Individual *Montipora capitata* colonies were tagged during the 2015 bleaching event in Kāne‘ohe Bay, Hawai‘i (Matsuda et al. 2019). These tagged corals represent pairs of colonies immediately adjacent to each other where one coral completely bleached while the other remained visibly healthy (Figure S1B). These tags have been actively maintained *in situ*, forming a biological library of known bleaching phenotypes during a natural thermal stress event.

We collected five replicate fragments from each of ten distinct genotypes (Drury, in review) from Reef 13 in Kāne‘ohe Bay in May 2019. These genotypes were evenly split between five historically bleached colonies and five historically non-bleached colonies selected from the pairs mentioned above.

We returned the coral fragments to the Hawai‘i Institute of Marine Biology, mounted them on aragonite plugs and acclimated them for four weeks in indoor mesocosms (hereafter ‘lab corals’) at ambient temperature conditions and a maximum midday irradiance of ∼400 μmol/sec. After acclimation, we subjected the lab corals to a five-day temperature stress at 31.4°C or a control (ambient) temperature of 28.4°C. We sampled each fragment before and after six days of heat stress with a 5 mm diameter dermal curette. Each sample was extracted in 500 μL of 70% methanol, kept on ice for 30 min and then stored at -80°C until shipping. Methanol extractions were shipped on dry ice to Michigan State University for untargeted metabolomics analysis.

### Mass spectrometry data collection and processing

Methanolic extracts were analyzed on a Thermo™ QExactive™ mass spectrometer coupled to a Vanquish Ultra High-Performance Liquid Chromatography (UHPLC) system. The mobile phase was 0.1% formic acid in Milli-Q water (channel A) and acetonitrile (channel B). The stationary phase was a reverse phase column Waters® Acquity® (Wood Dale, IL, USA) UPLC BEH C-18 column, 2.1 mm × 100 mm. The chromatographic runs were 12 min-long with linear gradients as follows: 0–1 min 2% B, 1–8 min 2–100% B. This 100% B solution was then held for 2 min followed by a switch to 2% B for the remaining 2 min. The injection volume was 10 µL, the flow rate 0.40 mL/min and the column temperature 60°C. Full MS^1^ survey scans and MS^2^ mass spectra for five precursor ions per survey scan were collected using electrospray ionization in positive mode with a scan range set from *m*/*z* 100 to 1500 for the full MS mode (minutes 1–10 of run). Raw files (.raw) were converted to .mzXML format for analysis. All files were processed with MZmine 2.53 software, GNPS and SIRIUS (Dührkop et al. 2019; Pluskal et al. 2010; Wang et al. 2016). Feature extraction for MS^1^ and MS^2^ was performed for a centroid mass detector with a signal threshold of 5.0 × 10^4^ counts. Chromatogram builder was run considering a minimum height of 1.0 × 10^5^ and a *m*/*z* tolerance of 10 ppm. Chromatograms were deconvoluted with a peak duration range of 0.0 to 3.00 min and a baseline cut-off algorithm of 1.0 × 10^5^. Isotopic peaks were grouped with a *m*/*z* tolerance of 0.02 Da and a retention time percentage of 0.05. Detected peaks were aligned through Join Aligner Module considering 0.02 Da and retention time tolerance of 0.02 min. The resulting peak list was gap filled, taking into account an intensity tolerance value of 0.001, 0.02 Da and retention time tolerance of 0.02 min. The data was converted to .mgf format and a feature quantification table was generated for running FBMN workflow on GNPS (Felix Nothias et al. 2019; Martin et al. 2019; Wang et al. 2016). FBMN was performed with a parent and fragment mass ion tolerance of 0.02 Da a cosine score of 0.65 and a minimum matched peaks minimum of 4. Feature-based molecular networking job is available at: https://gnps.ucsd.edu/ProteoSAFe/status.jsp?task=bb9b6126118c4ba1881f4483560012d3 and raw files are available in MASSIVE as <u>MSV000085272</u>.

### Molecular network analysis

Each node in a molecular network was assigned to a molecular family based on the presence of nodes with GNPS annotations and putative structures in the network (network annotation propagation). These annotations were visually inspected for characteristics of each molecular family. If no known metabolites were present, the network was considered as an unknown molecular family.

Meta-mass shift chemical profiling (MeMSChem) was used to analyze mass differences between related compounds indicative of biochemical transformations. We specifically focused on H2 additions to indicate the saturation state of lipid fatty acid chains. MeMSChem was performed according to the methods of Hartmann et al. 2017 and Quinn et al. 2020. Briefly, a reference node in the molecular network was used to determine the gain or loss of a mass shift from a related node connected through the cosine score. All nodes that qualified as having gained or lost the mass shift of interest were identified in the metabolomic feature table and the area-under-curve abundance summed for each coral sample. The total sum of these mass gains or losses were then compared for each coral sample between bleaching history phenotypes to identify the total abundance of metabolites becoming more or less saturated.

### Quantitative polymerase chain reaction (qPCR)

DNA was extracted from tissue samples taken from lab and validation corals with a CTAB-chloroform protocol (dx.doi.org/10.17504/protocols.io.dyq7vv). To determine the concentrations of different symbiont types and host cells in these samples, we used qPCR on an Applied Biosystems StepOnePlus system with two technical replicates. Actin assays were used to quantify symbionts of the genera *Cladocopium* and *Durusdinium*, and Pax-C assays were used to quantify the coral host cells (Cunning et al. 2015; Cunning and Baker 2013). Cycle threshold (CT) values were corrected for fluorescence and copy number using the equations of Cunning, Ritson-Williams, and Gates (2016). Samples were re-extracted and/or re-analyzed, if CT replicate standard deviation was greater than one between replicates or if only a single replicate amplified.

### Statistical analyses

Metabolite data was used to predict historical bleaching phenotype in lab corals using neural net classification (*JMP 14 Pro*, SAS Software) with a holdback proportion of 0.333, a learning rate of 0.1, a squared penalty method, and no random seed. We documented 100% accuracy when using all compounds in the dataset (including unannotated compounds). We then filtered the dataset to only known compounds and reclassified the data, again finding 100% accuracy. To identify which compounds were most important for distinguishing the historical bleaching phenotypes, we used supervised random forests classification analysis (*randomForest* R package) with 5000 trees to generate variable importance plots (VIPs) of all the compounds and the known compounds separately.

We examined the abundance of all molecules in the study using Principal Component Analysis (PCA) (*JMP 14 Pro*, SAS Software), then compared the distribution of genotypes and phenotypes using PERMANOVA in the R package *vegan* using 999 permutations. PCAs and PERMANOVAs were conducted independently for initial and validation sets using a) all compounds and b) known compounds. We compared the abundance of metabolites of interest and H2 gain (from mass change analysis) using t-tests. If data did not meet assumptions, we used Wilcoxon tests. We examined the abundance of highly informative lipids thought to be associated with symbiont genera (Rosset et al. 2019) using linear models. We calculated diversity metrics using the R package *vegan*.

We calculated the Normalized Heat Response (NHR) from the heat-stress time-series as the difference in metabolite abundance between final timepoints in the high temperature and control treatments less the initial difference using the formula below, where Ab = abundance

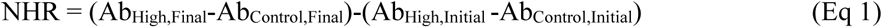

This calculation was conducted independently for each genotype and averaged across each phenotype, producing a Normalized Heat Response (i.e., normalized relative change in abundance due to temperature stress, corrected for change in the control), which integrates direction and magnitude of change. We defined outliers as molecules outside the first or third quartile ⨦1.5 IQR. Molecules outside of this range for one phenotype, but not the other, were considered phenotype-specific responses. Molecules within the threshold for both phenotypes were considered to have no response to heat stress. Molecules that responded in both phenotypes were classified as either Antagonistic (increasing in abundance in one phenotype and decreasing in the other) or Cooperative (increasing or decreasing in abundance in both phenotypes). We tested for independent proportions of outliers in predictive or all compounds using Fisher’s exact test.

We fit a linear model to NHR of each phenotype and tested the model against a linear model with a slope offset to examine differences from a slope of 1. We then examined variance in NHR for each phenotype using Levene’s test. To determine if molecular families respond to temperature stress, we analyzed each family-phenotype combination independently using a one-sample t-test to compare NHR to 0 with significance values adjusted by the Benjamini-Hochberg method (fdr < 0.05). Families with significant adjusted p-values for each phenotype were considered responsive to temperature stress. Several families only changed significantly in the bleaching phenotype and were designated as phenotype-specific heat responses. To examine if certain families had significantly higher magnitude NHR, we calculated distance from the origin for each molecule, classified each point by molecular family, and rank transformed the distances. We independently compared the ranks of each family to all other molecules in the study regardless of family assignment using a one-tailed Mann-Whitney U test and corrected p-values as above. We conducted enrichment analysis comparing the distribution of each family in the top 5% of NHR magnitude to expected based on the overall distribution using a Fisher’s exact test.

To examine the holobiont-wide response to thermal stress, we conducted a PCA on all experimental stress samples. To calculate change due to thermal stress, we used a two-way PERMANOVA with historical bleaching phenotype and timepoint (initial, final) as factors. We analyzed all molecules and then used the same framework to select specific molecular families to reanalyze: membrane lipids (betaines, phosphocholines), putative bioactive molecules (steroids, prostaglandins, eicosanoids, phosphatidic acids, PAF, Lyso-PAF) and microbial metabolites (microbial natural products, indoles).

### Validation data

To examine whether the phenotypic differences resolved in the pre-stress lab samples were consistent with coral samples taken directly from the reef, we returned to the original reef and collected a fragment from an additional 12 coral colonies with known bleaching histories that were not included in the original dataset (hereafter ‘validation corals’). Validation samples were collected in October 2019. These samples represent 12 distinct genotypes and were evenly split between historical bleaching phenotypes, serving as an *in situ* test-set with known phenotypes from the same tagging efforts in 2015. Coral fragments were sampled immediately, shipped, and analyzed by LC-MS/MS as described above. We applied the same previously described neural net classification and random forests analysis independently to the validation corals.

## Results

### Metabolomic signatures of historical bleaching phenotype

The LC-MS/MS metabolomic data produced 3,574 unique metabolite features of which 138 had a spectral match to a known compound in the GNPS libraries (level two according to the metabolomics standards initiative (Sumner et al. 2007)). All measures of metabolomic diversity including richness, Shannon entropy, and evenness were significantly higher in historically non-bleached corals (Wilcoxon test richness: p = 0.021, entropy: p = 0.010, evenness: p = 0.013; Figure S2). When using the total metabolome, there was strong differentiation between historically bleached and non-bleached corals (PERMANOVA bleaching history F = 11.03, R^2^ = 0.19, p < 0.001). To verify that discrimination between historical bleaching phenotypes was reproducible for corals sampled directly from the reef and between LC-MS/MS runs, we sampled a set of 12 additional, *in situ* coral colonies with known bleaching history that were not included in the original dataset (hereafter ‘validation corals’). The validation corals also had a strong metabolomic signature of the past bleaching event with a significant difference in the historical phenotypes when using the total metabolome (PERMANOVA bleaching history F = 4.18, R^2^ = 0.29, p < 0.001). To visualize this differentiation in phenotype we constructed two-dimensional principal component plots (Figure 1 A and B), which show clear clustering of the historically bleached and unbleached phenotypes with sub-clustering of genotypes nested within these phenotypic groups (PERMANOVA genotype F = 2.50, R^2^ = 0.051, p = 0.029). Furthermore, when using the total metabolome for neural net classification analysis, we were able to predict the historical bleaching phenotype with 100% accuracy in both the initial laboratory corals and the *in situ* validation corals (Figure S3).

**Figure 1.**
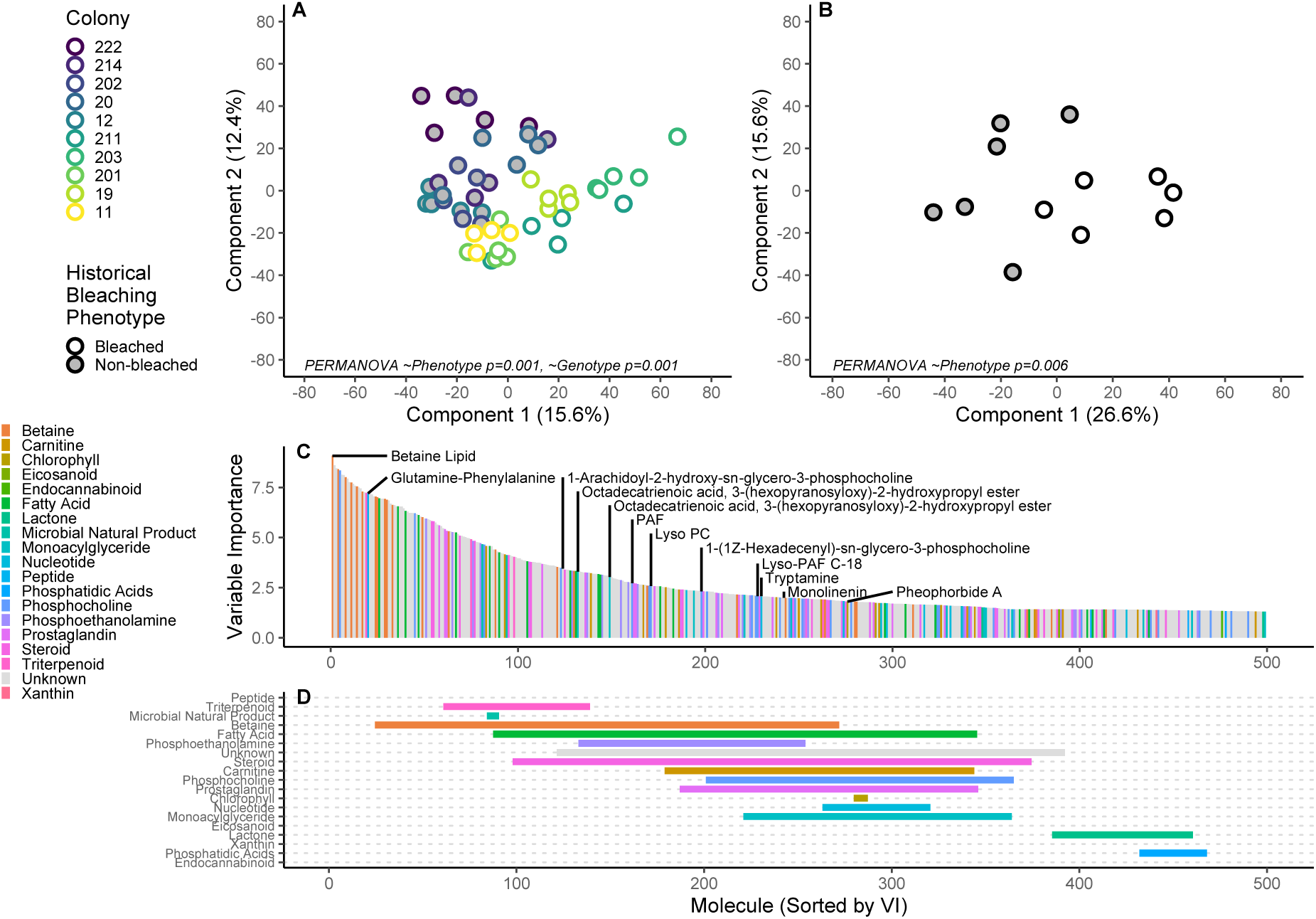
(A) PCA of lab corals using all compounds. Points (n = 5 per genotype) represent individual samples colored by genotype (n = 10). Fill represents historical bleaching phenotype (gray = non-bleached, white = bleached). (B) PCA of *in situ* validation set using all compounds. Points represent single samples from individual genotypes in the validation set which were not evaluated in panel A (n = 12). Fill represents historical bleaching phenotype (gray = non-bleached, white = bleached). (C) Top 500 most important compounds for discriminating between historical bleaching phenotypes, ranked by descending variable importance. Colors represent molecular families assigned to individual molecules. Labels denote molecules of biological interest and the most informative molecule for discriminating between phenotypes, a betaine lipid. (D) Relative variable importance of molecular families in top 500 most important compounds, arranged by ascending median rank (lower is more informative). Individual points correspond to ranked molecules in panel C, with horizontal bars showing first and third quartiles of each molecular family. Colors correspond to molecular families in panel C.

Supervised random forests classification analysis on bleaching history was used to identify metabolites that most strongly distinguished the two groups. Except for a Glu-Phe peptide, the top 100 strongest classifiers were all unknown molecules with no hits in the GNPS database. Thus, we used network annotation propagation from GNPS to classify unknowns into putative molecular families by their molecular network connection to known compounds (Figure 1C and D). This analysis revealed that small peptides, betaine lipids (see below), triterpenoids, microbial natural products, fatty acids, phosphoethanolamines, and steroids were, on average, the most important classes of molecules for distinguishing historical bleaching phenotypes (Figure 1D).

### Betaine lipids distinguish bleaching history

The strongest metabolite driver of coral bleaching history from the random forests classification was an unknown compound (*m/z*484.3272) containing signatures of choline in its MS/MS pattern and a retention time indicating significant hydrophobicity (6.4 min). This molecule was connected to a large molecular family with very similar MS/MS patterns, including a highly abundant compound with a parent mass *m/z*490.3688 (17th in the VIP; Figure 1C). Due to its abundance, we further investigated this metabolite using positive- and negative-mode MS/MS analysis. Its calculated molecular formula and fragmentation pattern revealed a C26H52NO7 lipid molecule with a choline head group and a C16:0 fatty acid ester (Figures 2A and S4). The molecular formula and subsequent MS/MS analysis identified this metabolite as a betaine lipid: diacylglyceryl-carboxyhydroxymethylcholine (DGCC 16:0/0:0). Identification of this compound led to annotation of others in the network including the primary driver described above (DGCC 16:3/0:0). Both of these molecules were highly enriched in non-bleached corals in both the lab corals (p < 0.001; Figure 2B and C) and the *in situ* validation set (p < 0.002; Figure S5A). Further analysis of this molecular network revealed that other betaine lipids with fully saturated fatty acid chains (DGCCs 14:0/0:0, 17:0/0:0, 20:0/0:0) were also strong classifiers (ranked in the top 0.08% of VIP predictors) and enriched in non-bleached corals (Figures 2A and S6). In contrast, long, highly unsaturated and diacyl forms of these compounds were significantly more abundant in bleached corals (e.g., *m/z*562.3685, DGCC 22:6/0:0, *m/z*822.5840, DGCC 40:9 (20:5/20:4); Figures 2D, E and S7). Six nodes were identified in the betaine lipid network as having parent masses above *m/z*700. Accurate mass analysis indicated all of these were diacyl forms of betaine lipids and were significantly more abundant in bleached corals (Table S1). The effect of bleaching history on the lipid saturation of these compounds was even found between two betaine lipids that were different only by a single fatty acid desaturation. The fully saturated lipid DGCC 18:0/0:0 (*m/z*518.4040) was significantly higher in non-bleached corals (p = 0.004; Figure 2F), whereas its monounsaturated relative DGCC 18:1/0:0 (*m/z*516.3845) was not (p = 0.848; Figure 2G). This finding led to further investigation of saturation properties throughout the metabolomic dataset (see below).

**Figure 2.**
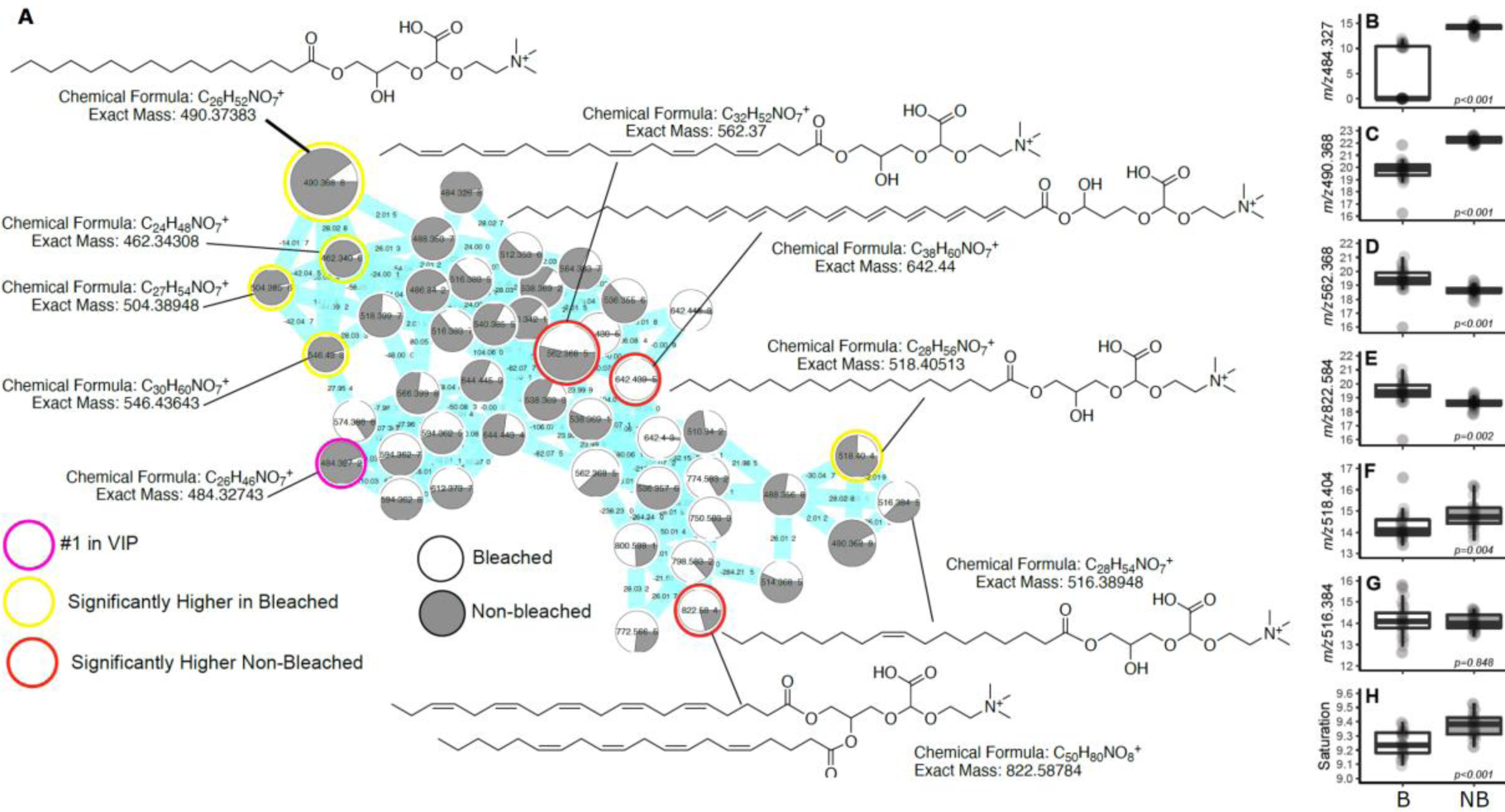
Molecular network and boxplots of the betaine lipid molecular family. Each node in the network (A) represents a unique MS/MS spectrum (i.e., a unique metabolite) and is labelled by the parent mass of the compound. The gray-white pie graph in each node is derived from the abundance of each metabolite in either bleached (white) or non-bleached (gray) corals. Boxplots B) through G) show the log area-under-curve abundances in the two bleaching phenotypes of the highlighted lipids. (H) Shows the saturation state of all metabolites in the molecular network. The colored circles highlight the nodes that are featured by their structure and/or boxplots. Boxplots are median with quartiles and whiskers extending 1.5 IQR beyond quartiles. Putative structures of the compounds are also shown along with molecular formulas and exact calculated masses, the exact positions of the fatty acid desaturations are uncertain.

We then investigated the relationships between highly abundant representatives of the saturated and unsaturated betaine lipids and the ratio of *Durusdinium* and *Cladocopium* algal symbionts to coral host cells. There was a significant positive correlation between the abundance of a long, unsaturated betaine lipid (*m/z*562.3685, DGCC 22:6/0:0) and the density of the heat-sensitive *Cladocopium* algal symbionts (p < 0.0001, R^2^ = 0.325), and a significantly negative correlation between a shorter, fully saturated version of the lipid (DGCC 16:0/0:0, *m/z*490.3688, p = 0.0047, R^2^ = 0.167; Figure S8A and B). There was no significant correlation between the heat-resistant *Durusdinium* algal density and these lipids (Figure S8C and D).

### Edge analysis to infer chemical changes

To further analyze the total metabolome, meta-mass shift chemical (MeMSChem) profiling (Hartmann et al. 2017) was used to characterize chemical modifications in the molecular network. MeMSChem analysis revealed that there were significantly higher H2 additions in historically non-bleached corals in both the lab (t-test p < 0.001, Figure 2H) and validation set (t-test p = 0.029, Figure S5G) indicating that lipids in the non-bleached corals were significantly more saturated.

### Analysis of known compounds

When using only the known compounds, neural net classification was also able to distinguish between historical bleaching phenotypes with 100% accuracy. This was visualized in two-dimensional principal coordinates plots in Figure 3A and B, which show that the samples separate significantly by phenotype (Lab corals: PERMANOVA ∼phenotype p < 0.001, Validation corals: PERMANOVA ∼phenotype p = 0.004). Random forests analysis was used to generate variable importance scores to rank known compounds in order of importance for predicting phenotype (Figure 3B). The most important known compound for predicting phenotype was a glutamine-phenylalanine dipeptide, which was significantly enriched in historically non-bleached corals (p < 0.0001; Figure S9A). A majority of the important predictors were phosphocholine derivatives and monoacylglycerides (Figure 3B and C). Several classes of bioactive lipids, including steroids and prostaglandins, also demonstrated high predictive power. Platelet Activating Factor (PAF), a host-derived bioactive lipid, was significantly enriched in historically bleached corals (p < 0.0001; Figure S9B). The inactive precursor of PAF, Lyso-PAF, also discriminated between phenotypes, but was significantly higher in historically non-bleached corals (p < 0.0001; Figure S9C). Pheophorbide A, a zooxanthellae-produced chlorophyll break-down product, was significantly enriched in historically non-bleached corals (p < 0.0001; Figures 3C and S9D), as was the photosynthetic compound fucoxanthin (p = 0.0008; Figure S9E). In the *in situ* validation corals, all compounds of interest (Glu-Phe, Lyso-PAF, pheophorbide A, and fucoxanthin), except PAF, followed the same trend as they did in the lab corals (Figure S9 F-J).

**Figure 3.**
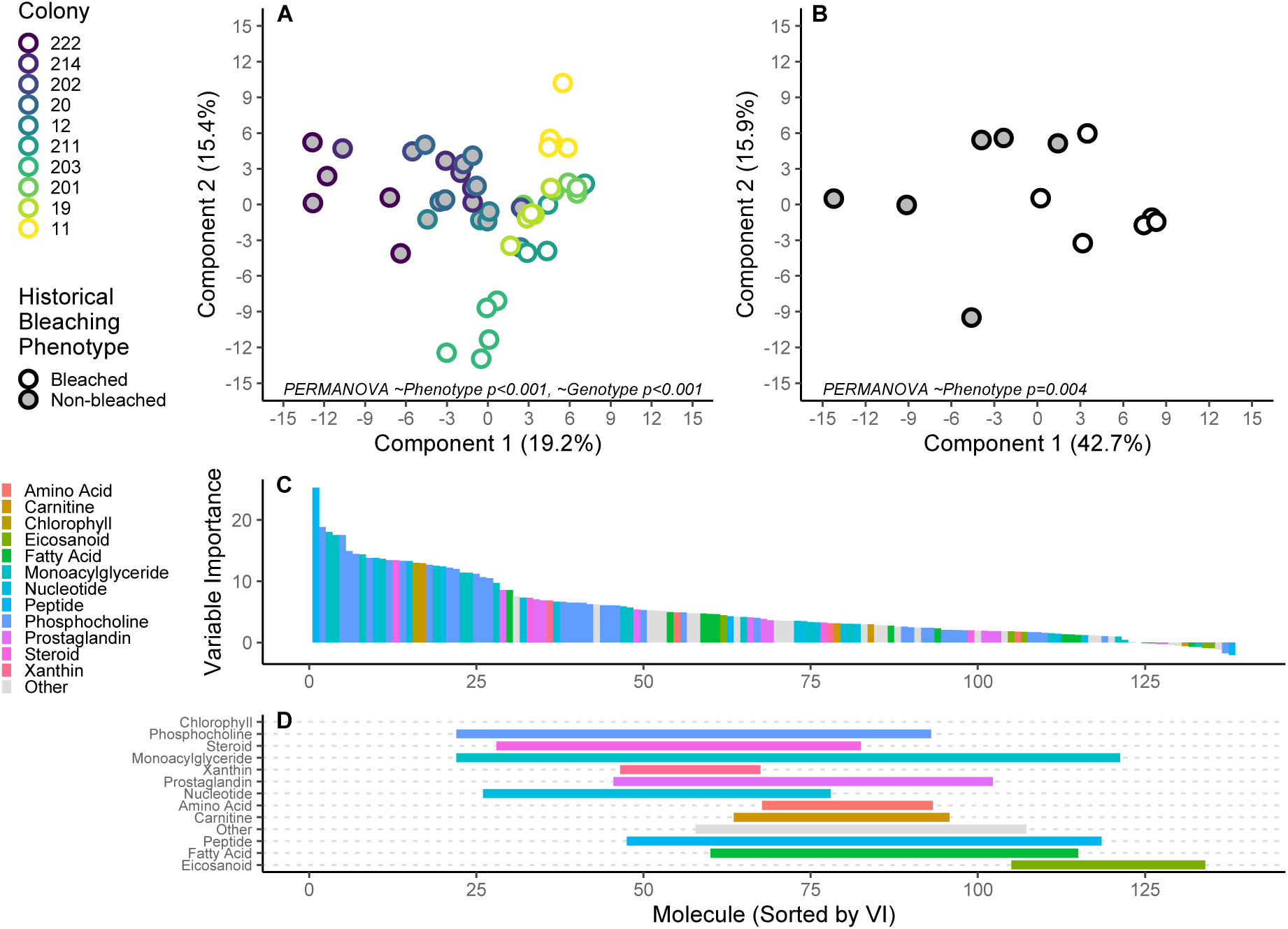
(A) PCA of lab corals using only known compounds. Points (n=5 per genotype) represent individual samples colored by genotype (n=10). Fill represents historical bleaching phenotype (gray = non-bleached, white = bleached). (B) PCA of field validation set using only known compounds. Points represent single samples from individual genotypes in the validation set which were not evaluated in panel A (n=12). Fill represents historical bleaching phenotype (gray = non-bleached, white = bleached). (C) All known compounds ranked by descending variable importance for discriminating between historical bleaching phenotypes. Colors represent molecular families assigned to individual molecules corresponding to Figure 1. (D) Relative variable importance of molecular families in all known compounds, arranged by ascending median rank (lower is more informative). Individual points correspond to ranked molecules in panel C, with horizontal bars showing first and third quartiles of each molecular family. Colors represent molecular families as in panel C.

Linear regression analysis demonstrated that the host-derived bioactive lipids Lyso-PAF and PAF had significant relationships (p ≤ 0.0005) with the symbiont community and symbiont-derived metabolites (Figure S10). Specifically, the Lyso-PAF:PAF ratio had significant positive correlations with the *Durisdinium* to *Cladocopium* ratio (R^2^ = 0.366, p = 0.0005), pheophorbide A (R^2^ = 0.325, p < 0.0001), and the saturated betaine lipids *m/z*484.3272 (R^2^ = 0.763, p < 0.0001) and *m/z*490.3688 (R^2^ = 0.631, p < 0.0001). Conversely, there was a significant negative correlation with the highly unsaturated betaine lipid *m/z*562.3685 (R^2^ = 0.366, p < 0.0001).

### Experimental temperature stress

After acclimation and initial sampling, the lab corals were subjected to a six-day heat stress at 31.4 ± 0.08°C (mean ± 1SD) or control (ambient) at 28.4 ± 0.15°C temperature profile. Each fragment was sampled before and at the conclusion of the six-day heat stress treatment.

There was a significant positive correlation between the Normalized Heat Response (NHR) in the two phenotypes (p < 0.001) with substantial explanatory power (R^2^ = 0.2409) and a slope significantly less than 1 (m = 0.565, p < 0.001). Variance in the non-bleached NHR was 32.5% higher than in bleached NHR but was not significantly different (p = 0.161). Mean NHR was significantly larger in non-bleached corals (p = 0.005).

Using NHR of all molecules, 1014 compounds (28.4%) had outlier magnitudes of change, including 34 of the 100 most important compounds for discriminating between historical bleaching phenotypes in the pre-stress lab corals (Figure 4A). Thus, most of the compounds (66 out of 100) that were important for distinguishing between the initial bleaching history phenotypes did not change substantially under heat stress (Figure 4B). Most of the top discriminatory compounds that responded to heat stress (24 of 34) only changed in a single phenotype (i.e., these compounds only changed in the historically bleached (n = 12) or non-bleached corals (n = 12) under stress). Three molecules changed cooperatively (decreasing abundance in both phenotypes) and seven molecules had antagonistic changes in abundance, all of which increased in the non-bleached phenotype and decreased in the bleached phenotype.

**Figure 4.**
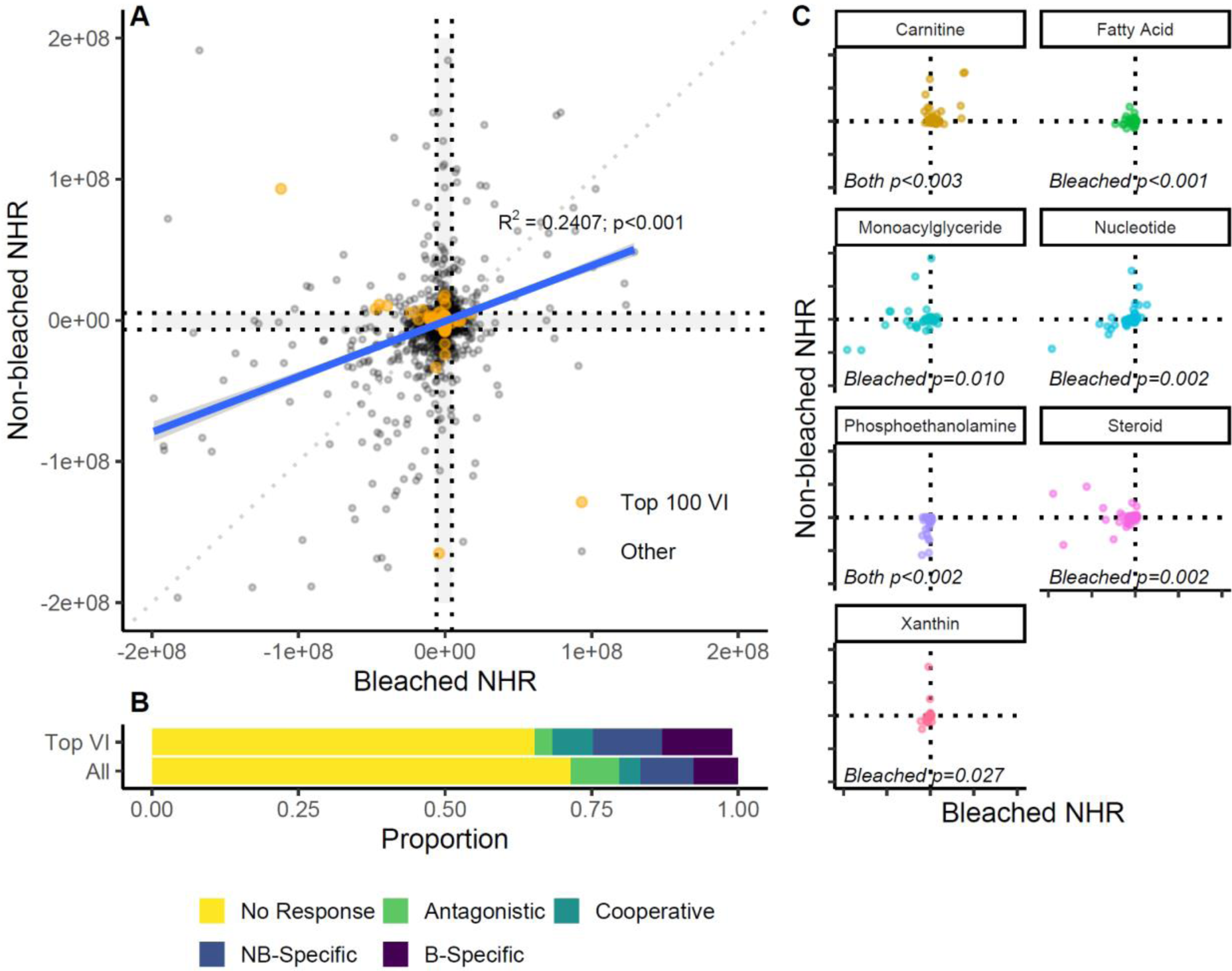
(A) Response of individual metabolites to temperature stress. Each point represents the Normalized Heat Response (NHR, control corrected change in abundance) of an individual compound. Orange points denote the top 100 variables for discriminating between phenotypes before bleaching stress. We created outlier cutoffs for each axis using the first or third quartile ⨦1.5 IQR as a threshold (dashed lines). Molecules that fell outside this range for one phenotype, but not the other, were considered phenotype-specific responses. Molecules that fell within the threshold for both phenotypes were considered no response to heat stress. Molecules that responded in both phenotypes were classified as either cooperative (increasing or decreasing in both phenotypes) or antagonistic (increasing in abundance in one phenotype and decreasing in the other). (B) Proportion of compounds by classification for top 100 VI and all compounds. NB stands for Non-Bleached and B stands for Bleached. (C) Family-specific changes due to temperature stress. All families shown were significantly different from zero NHR for at least one phenotype; significance patterns are inset in each figure (Bleached = bleached-specific, Both = significant change in both phenotypes). Molecular family colors correspond to Figures 1 and 3.

Analysis of family response to temperature stress (Figure 4C) found five families with significant bleached-specific NHR (nucleotides, steroids, xanthins, fatty acids, monoacylglycerides; p < 0.027), and two families with significant changes in both phenotypes (carnitines, phosphoethanolamines; p < 0.037; Table S2). Steroids, monoacylglycerides, endocannabinoids, indoles and chlorophyll had significantly larger magnitude NHR than the average molecule (p < 0.046; Table S3). Betaines, chlorophyll, indoles, monoacylglycerides and steroids were significantly enriched (p < 0.024) in molecules with the top 5% magnitude of change (Table S4).

There was a significantly different response to heat stress between phenotypes in the total metabolome (interaction p = 0.018; Figure 5A), contributing to a larger difference between phenotypes after thermal stress. In membrane lipids, there was a significant effect of phenotype (p = 0.002, Figure 5B), and all historically bleached corals converged due to response in a single direction. Bioactive molecules were also significantly different between phenotypes (p < 0.001, Figure 5C), and had a nearly orthogonal response to heat stress between phenotypes. Putative microbial products were significantly different between phenotypes (p < 0.001; Figure 5D) and displayed a highly variable response to temperature stress based on phenotype.

**Figure 5.**
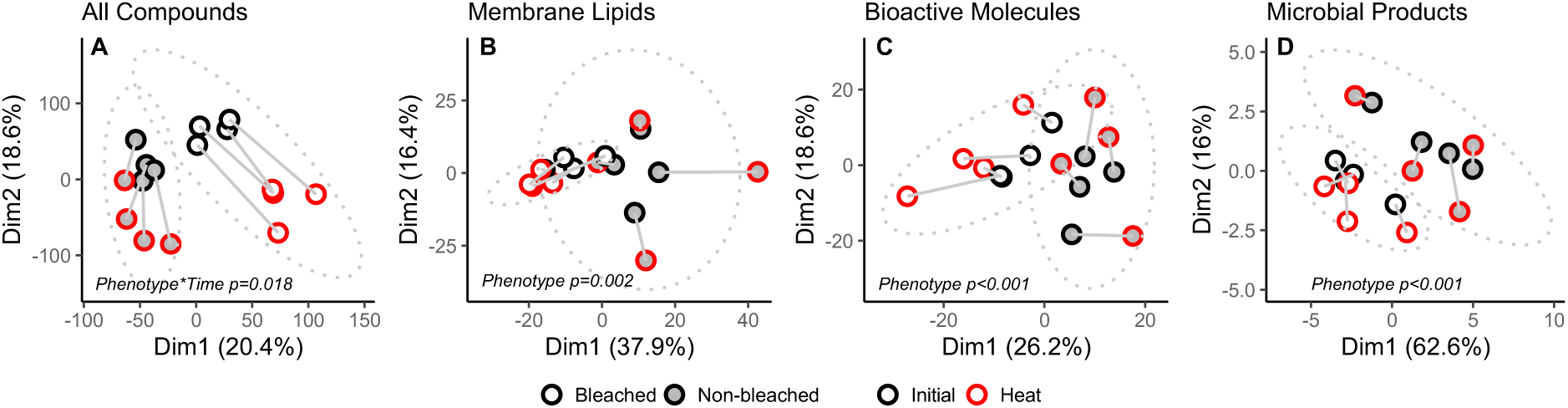
Multivariate response to heat stress for (A) the total metabolome, (B) membrane lipids, (C) bioactive molecules, (D) putative microbial products. Fill represents historical bleaching phenotype (gray = non-bleached, white = bleached). Color represents initial and heat-stress timepoints (black = initial, red = heat-stress). Gray ellipses represent 95% confidence interval of phenotypes. Gray bars connect paired before-after samples.

## Discussion

The data presented herein illustrate that there is a strong metabolomic signature of past bleaching history even when corals are not experiencing thermal stress. During temperature stress these inherent biochemical differences were further enhanced. This signature was simultaneously observed in the coral host, the symbiotic algae, and putative bacterial natural products. The major difference between bleaching phenotypes was a group of betaine lipids associated with the algal symbionts for which the saturated forms were significantly overrepresented in historically non-bleached corals. Conversely, the long chain and unsaturated forms were enriched in historically bleached corals. The host-derived bioactive lipid PAF and its precursor Lyso-PAF were also significantly different based on bleaching history and the ratio of these compounds significantly associated with the symbiont community and the betaine lipids driving bleaching history, indicating a possible host regulation of heat tolerant or heat sensitive symbionts. When corals were subjected to experimental temperature stress a majority of the compounds that distinguished historical bleaching phenotypes did not change substantially, and if they did, they typically changed in one phenotype or the other resulting in further differentiation of the two phenotypes.

### Betaine lipids from the symbiont

Top predictors for distinguishing historically bleached and non-bleached corals were novel, totally saturated betaine Lyso-lipid molecules. The major difference between these lipids in the historical bleaching phenotypes was that non-bleached corals had higher degrees of saturation than the historically bleached corals. A similar phenomenon was recently identified in Rosset et al. 2019, who showed that isolated heat-tolerant *Durusdinium trenchii* algae also had higher levels of saturation of betaine lipids, along with other lipid classes. Here, we report this in corals taken directly from the reef but also show that the major drivers of bleaching history in these native corals were fully saturated betaine lipids, those not reported in (Rosset et al. 2019). We have also shown that bleached corals contained far higher abundances of diacyl-betaine lipids, as opposed to the Lyso-lipid form. Diacyl lipids are more likely to form bilayers, indicating the non-bleached corals may have fundamentally different functions for this lipid class. It is therefore apparent that thermal tolerance of corals is tightly linked to the saturation and acylation of these, and likely other, lipid groups in both a laboratory setting and in response to historical bleaching events *in situ*. Though the exact location of these betaine lipids in the algae is unknown, they may contribute directly to the thermal stability of the plastids, a known requirement for thermal tolerance (Mansour et al. 2018), or they could provide thermal stability to other membranes of the symbiont. It is also possible that their role in thermal tolerance is through regulatory mechanisms, similar to other coral bioactive lipids (Quinn et al. 2016; Roach et al. in press). Regardless of their organelle source or mechanism of function, the enrichment of these lipids in resilient corals on a live reef provides potential for their use as biomarkers of coral resilience and further identifies betaine lipid saturation as an important biological mechanism behind coral bleaching.

### Host-produced metabolites

The bioactive lipid Platelet Activating Factor (PAF) and its precursor Lyso-PAF were significantly different between historical bleaching phenotypes and can be linked to the coral host (Quinn et al. 2016). These host-derived molecules are known to play a role in coral physiology and defense, with higher amounts of PAF being produced when corals are undergoing stress (Galtier d’Auriac et al. 2018). Interestingly, the activated form of the molecule (PAF) was higher in historically bleached corals, while the inactive form (Lyso-PAF) was higher in non-bleached corals, even when the corals were not undergoing temperature stress. These bioactive lipids are believed to play a role in coral self/non-self-recognition and immune response (Galtier d’Auriac et al. 2018; Quinn et al. 2016). Further support for this role comes from the strong association between the ratio of Lyso-PAF to PAF with the symbiont communities and their associated metabolites, including the algal-derived betaine lipids. This finding indicates interplay between the host immune system and the symbiont community in determining response to temperature stress and suggests that host immune modulation may contribute to the vastly different algal symbiont communities in adjacent corals. This link between self/non-self-recognition, immune response, and host symbiont choice provides an area rich for potential new research. These findings also further the discussion on the role of bioactive lipids in coral physiology as well as ancient links between human and coral immune responses (Quistad et al. 2014).

### Symbiotic algae-produced metabolites

The data also demonstrate signatures of past bleaching in other metabolites from the symbiotic algae in addition to the betaine lipids. In particular, the chlorophyll breakdown product pheophorbide A and the accessory pigment fucoxanthin were both significantly enriched in non-bleached corals. This trend was observed in the pre-stress corals but became even stronger during heat stress with a > 1,000X increase in both compounds for the non-bleached corals. This suggests that corals that do not bleach during temperature stress may be degrading chlorophyll and producing more photoprotective photosynthetic pigments such as fucoxanthin and other carotenoids. The breakdown of the light harvesting complex in chlorophyll *a* has previously been linked to bleaching resistance (Hill, Larkum, and Kramer 2012) as has the production of photoprotective xanthophylls (Venn et al. 2006). However, the stability of this pattern under non-thermally stressful conditions has not been shown before.

### Fatty acid saturation state

Edge analysis (Hartmann et al. 2017) of the total metabolomic networks demonstrated that the metabolome (which consisted mostly of lipids) of the non-bleached corals were significantly more saturated. This suggests that the lipids in the historically non-bleached corals are more thermally stable and more resistant to oxidative stress, which could provide a mechanism for corals to deal with the excess free radicals that can be produced through photosynthesis during times of temperature stress. Higher saturation could be particularly important for the thylakoid membranes of the zooxanthellae (Tchernov et al. 2004), as protecting them from oxidation may prevent triggering of the bleaching response. The combination of reducing free radical production by switching from chlorophylls to xanthophylls and having membranes that are more resistant to oxidative stress may underlie the tolerance of specific coral holobionts to temperature stress.

### Effects of experimental temperature stress on the coral metabolome

Many of the molecules that distinguish historical bleaching phenotypes do not respond to heat stress, suggesting that latent or causative effects of natural historical bleaching are partially decoupled from the active short-term heat response. These impacts likely relate to the different symbiont communities harbored within each phenotype, which greatly impact membrane lipids throughout our dataset.

Conversely, the total metabolome exhibits substantial change in response to heat stress that is more variable in historically bleached corals and further separates phenotypes from their pre-stressed states. Interestingly, heat-stressed non-bleached corals do not become more similar to their bleaching sensitive counterparts, suggesting that broadly different pathways define thermal tolerance. The strong response of individual molecular families to heat stress supports a phenotype-specific response that goes beyond symbiotic drivers and is illustrated by significant changes in diterpenoids, eicosanoids and fatty acids in one phenotype but not the other.

We also resolve significant differences between phenotypes in membrane lipids (likely due to symbionts), bioactive molecules, and putative microbial products in response to heat stress. While we are unable to quantify the relative contribution of each component of the holobiont to thermal stress, these data illustrate that natural bleaching susceptibility is a simultaneous product of the broad interaction between host cnidarian, algal symbionts, and bacterial metabolites. Further, the different components of the holobiont actively respond to thermal stress in a unique manner in each phenotype, suggesting positive feedback may make more resilient corals even stronger. Our data also show substantial genotypic influence nested within phenotypes in structure of the total metabolome and in response to temperature stress, suggesting a strong influence of the coral host on holobiont metabolism.

### Implications for coral reef restoration

Historically, marine conservation has focused on passive habitat protection to mitigate the effects of global stressors on coral reefs (Boström-Einarsson et al. 2020). However, the increasing frequency and severity of coral bleaching events necessitates active approaches to coral reef restoration, including asexual propagation, transplantation, larval enhancement, and assisted evolution. As marine heat waves and mass bleaching events become more common, it is important to ensure that reefs are restored using resilient coral stock (van Oppen et al. 2017; van Oppen et al. 2015); without this consideration the success rates of these efforts will likely be quite low (Baums et al. 2019). Thus, the identification of resilient corals for conservation and management has become critically important for the future of coral reefs but can be time-consuming as it often depends on natural bleaching outcomes or experimental results. Our metabolomics approach presents an opportunity for fast and cost-effective screening for resilient corals outside times of thermal stress. By screening for metabolite biomarkers, such as saturated betaine lipids or PAF, managers can test and select resilient coral stocks for restoration when corals are healthy. These selective restoration strategies are needed to restore reefs that will be resilient during future bleaching events (Bay et al. 2017), supporting durable, climate-wise coral ecosystems.

## Conclusion Statement

This study demonstrates a metabolomic signature associated with past bleaching in several components of the coral holobiont. This difference between phenotypes is largely driven by the saturation state of betaine lipids associated with the zooxanthellae and host immune response molecules. Overall, we find that structural membrane lipids, bioactive lipids, photosynthetic compounds, and microbial natural products all have distinct patterns associated with a coral’s past bleaching phenotype and response to heat stress. Additionally, this study shows that sublethal heat stress does not seem to alter this strong metabolomic signature of the two historical bleaching phenotypes, but rather amplifies the distinction between phenotypes. This work provides insight into the biochemical and physiological mechanisms involved in coral bleaching and symbiosis ecology and provides a novel tool for restoring resilient coral reefs in the face of global climate change.

## Acknowledgments

We dedicate this paper to Dr. Ruth Gates. We are thankful for her inspiration to use molecular tools to investigate corals and develop real-world solutions to the global coral reef crisis. We thank Nina Bean, Carlo Caruso, Luke Kikukawa, Shayle Matsuda, Josh Hancock, Ariana Huffmyer, Mariana Rocha de Souza, and the Gates Coral Lab for assistance with this project. This work was funded by the Paul G Allen Family Foundation. Coral collections were made under DLNR permit SAP 2020-25 to HIMB.

