## Supplemental Material for "Metabolomic signatures of coral bleaching history"

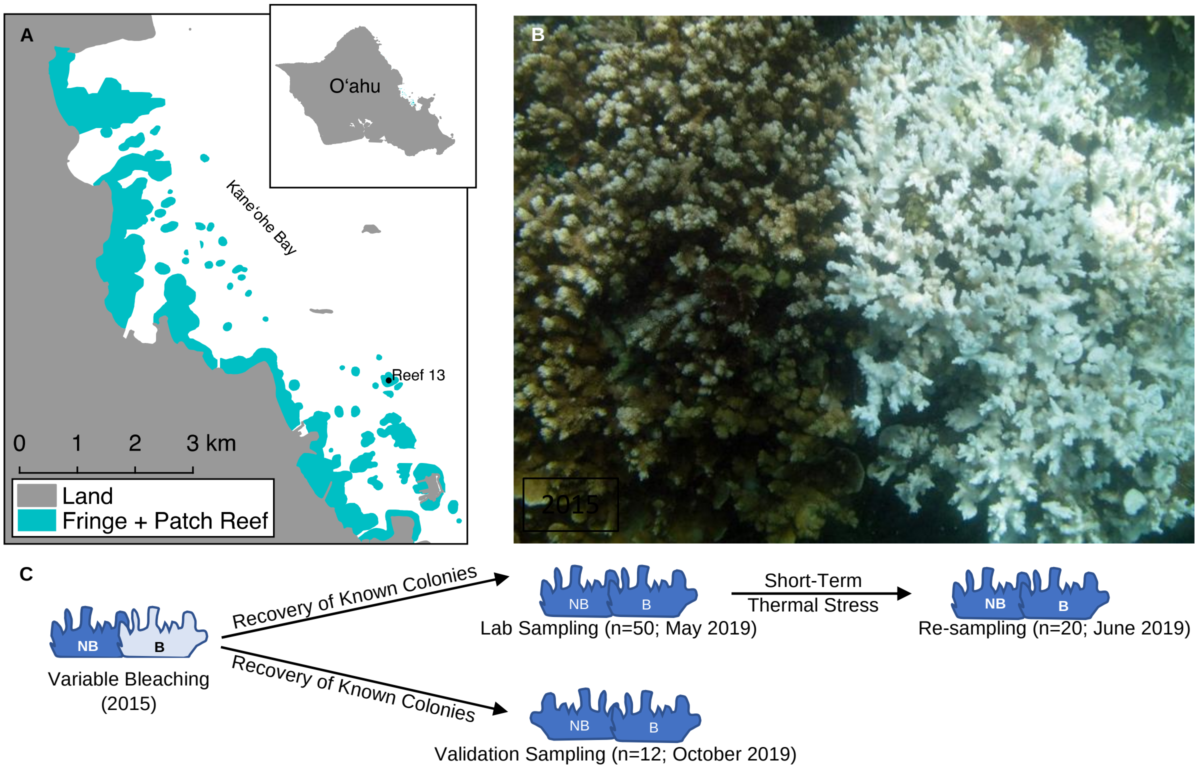


**Supplemental Figure 1.** Corals were collected from Reef 13 in in Kāne‘ohe Bay, O‘ahu, Hawai‘i (A). A representative pair of bleached and non-bleached corals at Reef 13 during the 2015 bleaching event (B). Overall sampling schematic (C). Abbreviations in panel C are as follows: NB = Non-Bleached and B = Bleached.

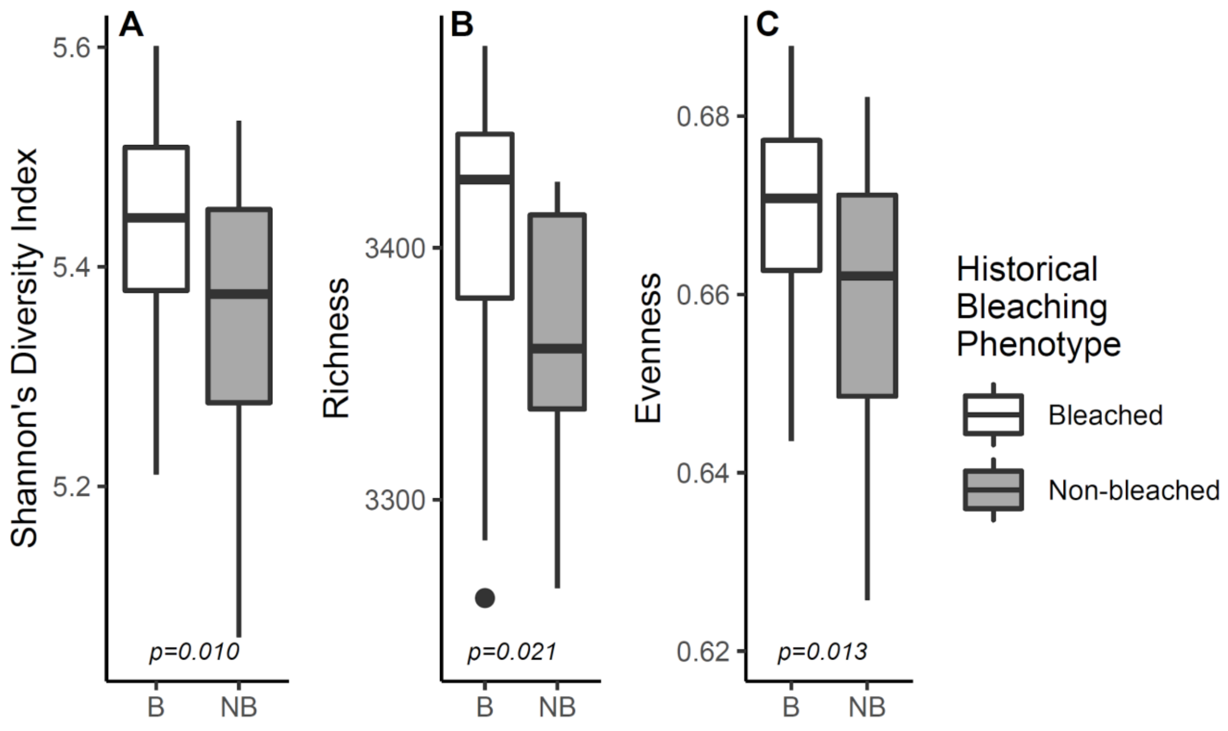


**Supplemental Figure 2.** Diversity metric for the metabolomes of bleached (B) and non-bleached (NB) corals including Shannon's entropy (A), richness (B), and evenness (C). Boxplots are median with quartiles and whiskers extending 1.5 IQR beyond quartiles.

**Lab Corals**

A.

| Actual BleachingHistoryPhenotype | Predicted Rate | |
|---|---|---|
| | Bleached | Non-bleached |
| Bleached | 1.000 | 0.000 |
| Non-bleached | 0.000 | 1.000 |

**Validation Corals**

B.

| Actual BleachingHistoryPhenotype | Predicted Rate | |
|---|---|---|
| | Bleached | Non-bleached |
| Bleached | 1.000 | 0.000 |
| Non-bleached | 0.000 | 1.000 |

**Supplemental Figure 3.** Neural net confusion matrices for predicting coral phenotypes in the 'lab corals' (A) and the 'validation corals' (B).

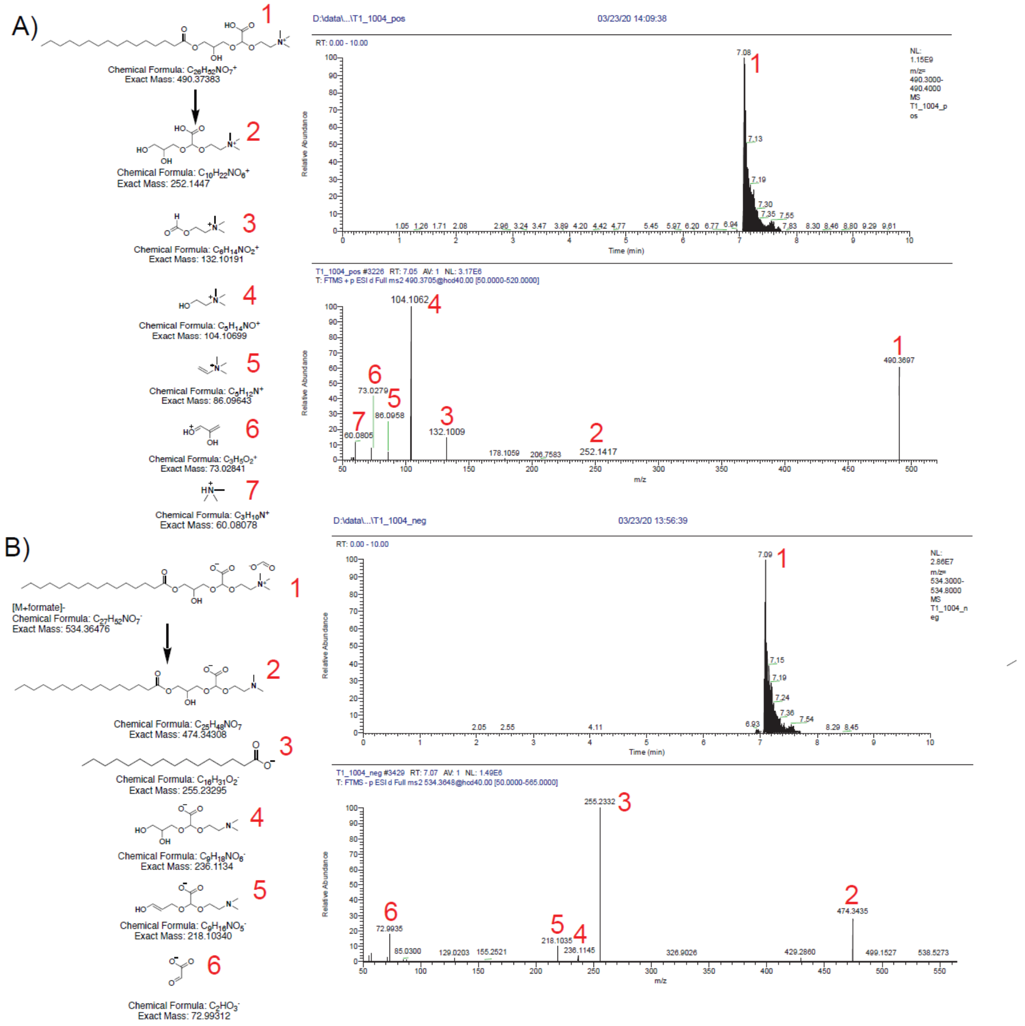

**Supplemental Figure 4.** Extracted ion chromatogram and MS/MS spectra of betaine lipid DGCC 16:0/0:0 described in this manuscript in positive (A) and negative (B) modes. The proposed annotation of each selected fragment ion is highlighted by a red number.

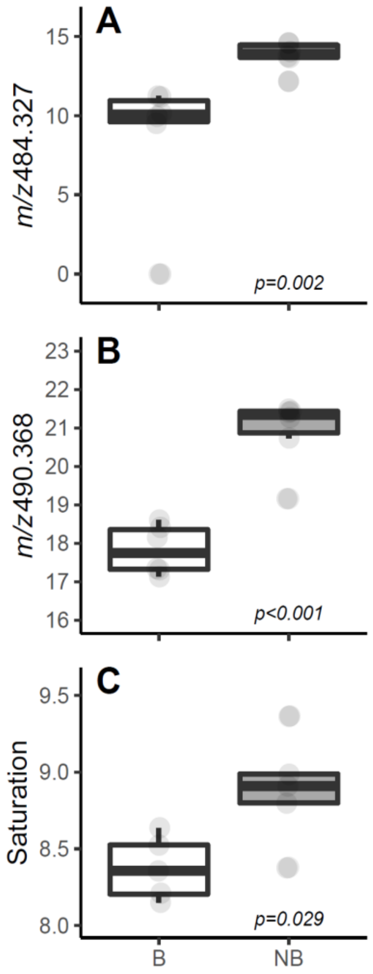


**Figure S5.** Box and whisker plots of A) the most important molecule for distinguishing between phenotypes, B) the most abundant fully saturated betaine lipid, and C) the saturation state of all molecules in the metabolome. Boxplots are median with quartiles and whiskers extending 1.5 IQR beyond quartiles.

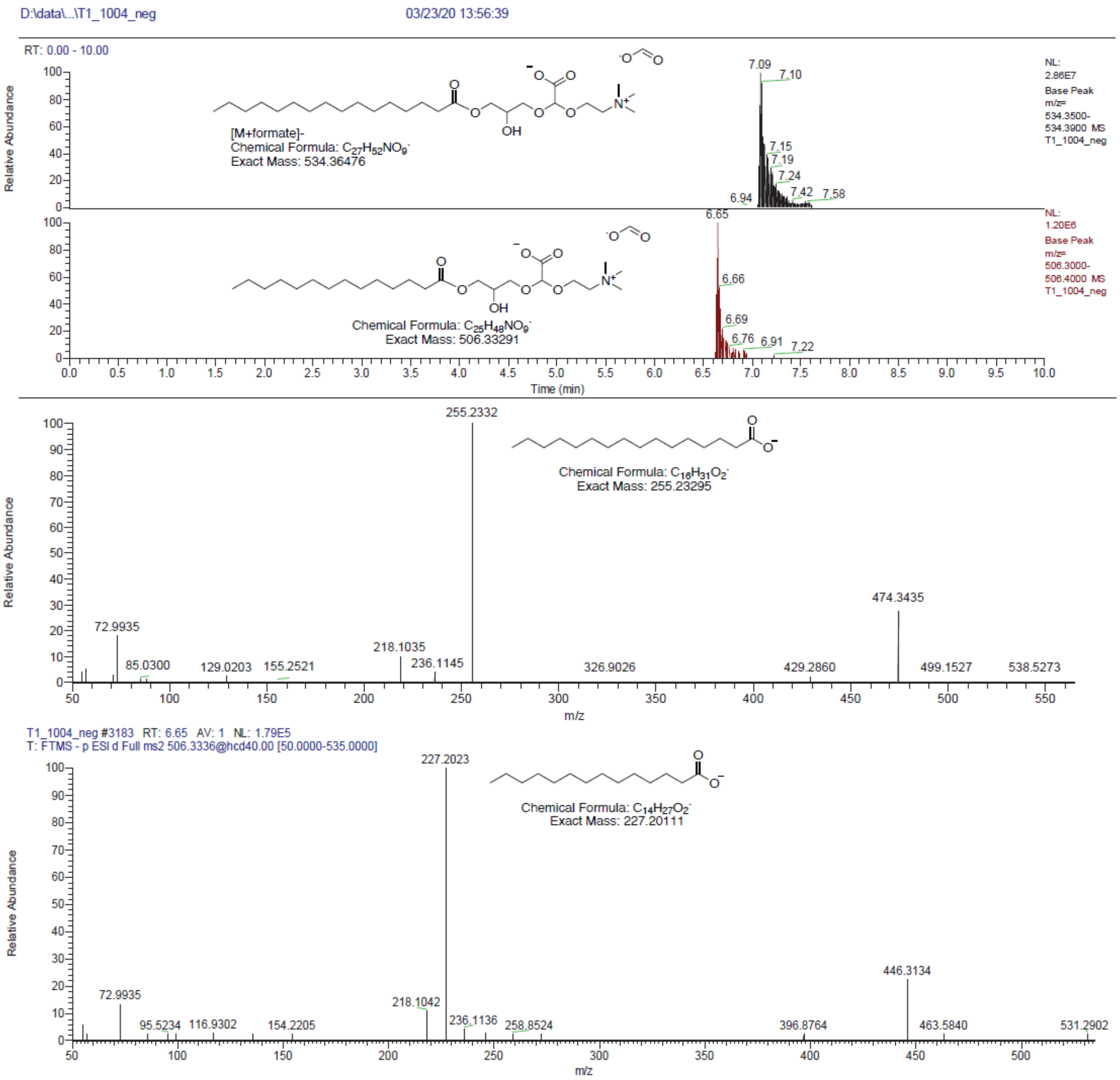


**Supplemental Figure 6.** Extracted ion chromatograms for [M+formate]$^-$ ions and MS/MS spectra of betaine lipids DGCC 16:0/0:0 and DGCC 14:0/0:0 with varied fatty acid chain lengths as determined using negative ion mode.

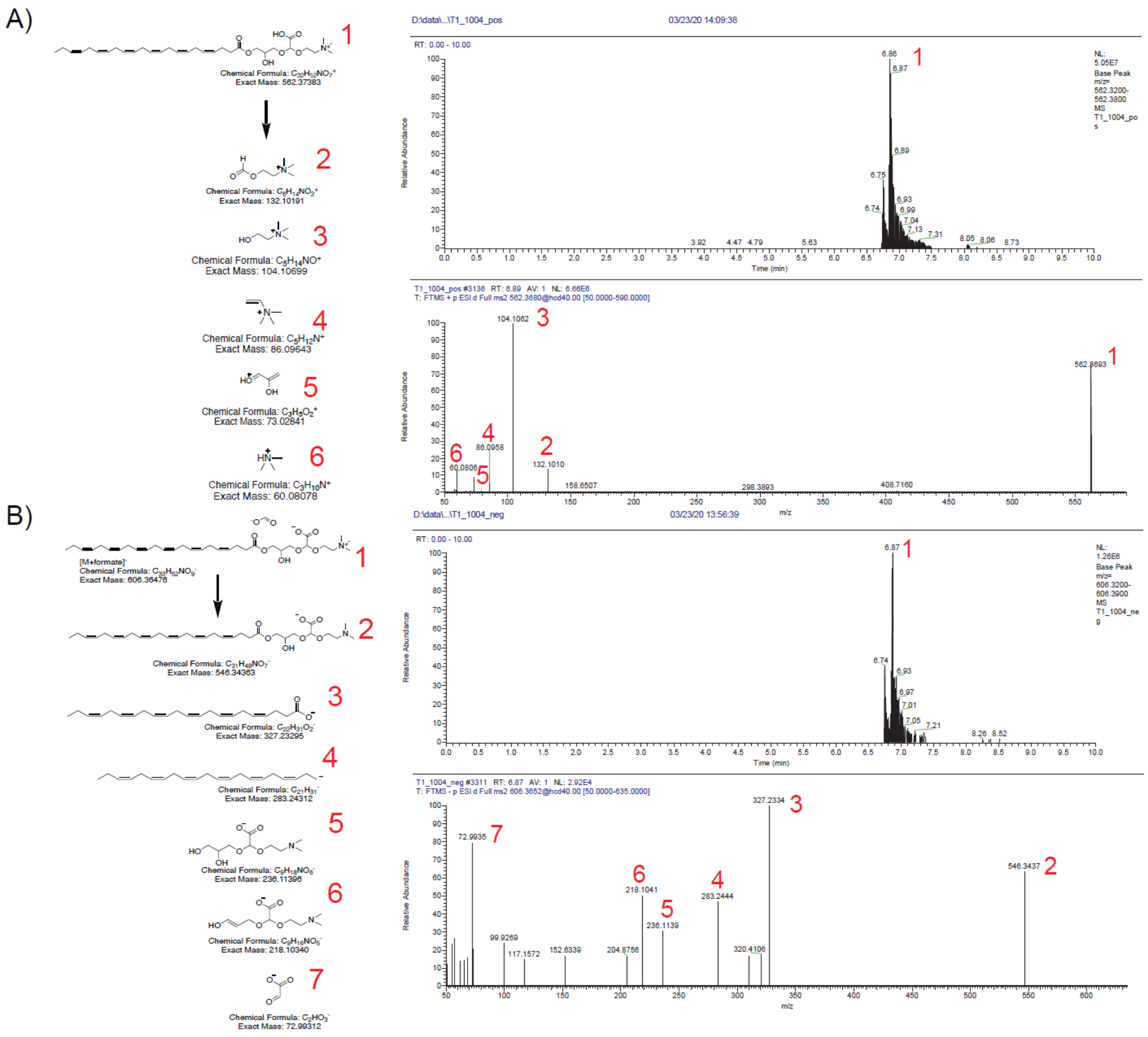


**Supplemental Figure 7.** Extracted ion chromatograms and MS/MS spectra of betaine lipid DGCC 22:6/0:0 described in this manuscript in positive (A) and negative (B) modes. The annotation of each selected fragment ion is highlighted by a red number.

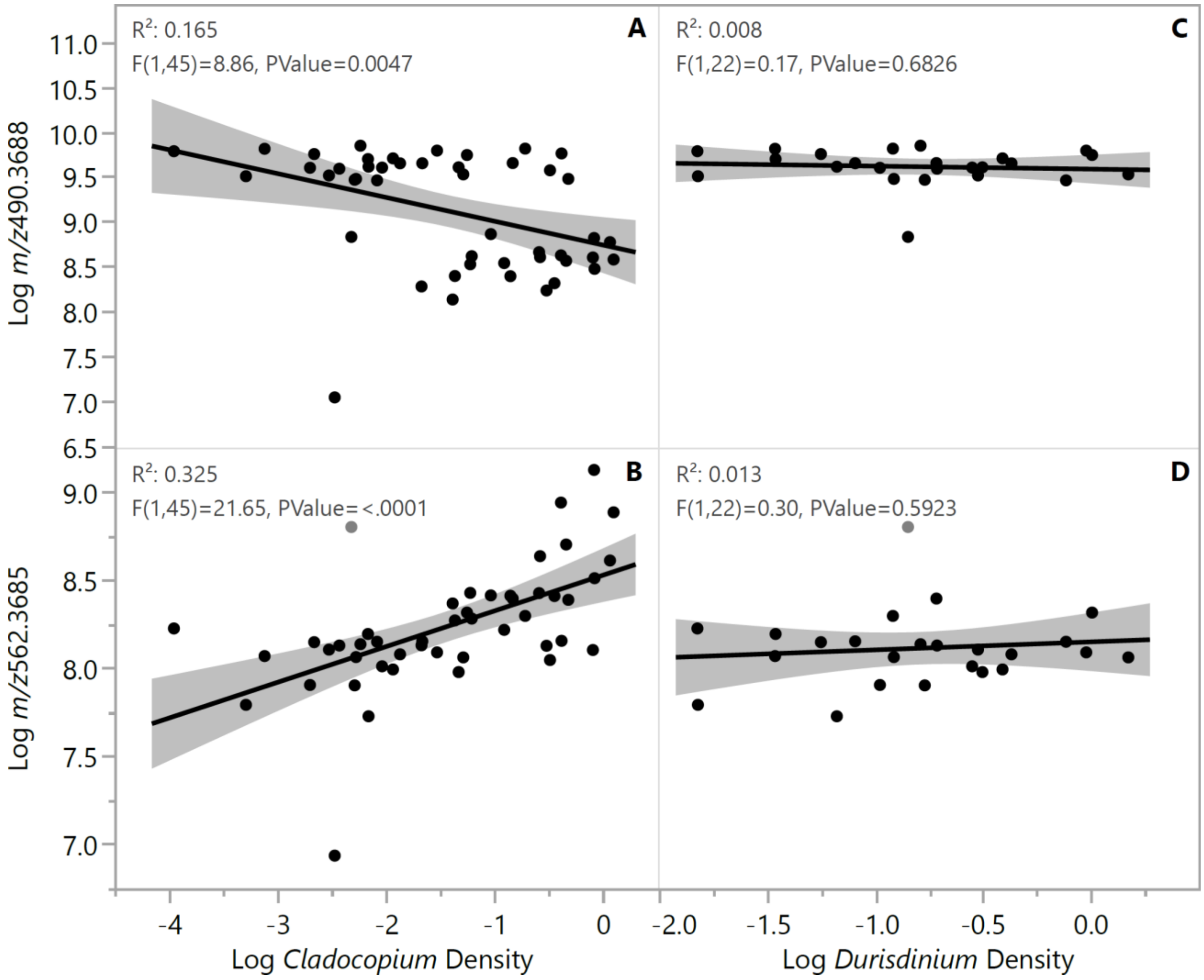


**Figure S8.** Linear regression of saturated (A,C) and unsaturated (B,D) betaine lipids versus *Cladocopium* (A,B) and *Durisdinium* (C,D) algal symbionts. Shaded areas represent 95% confidence intervals.

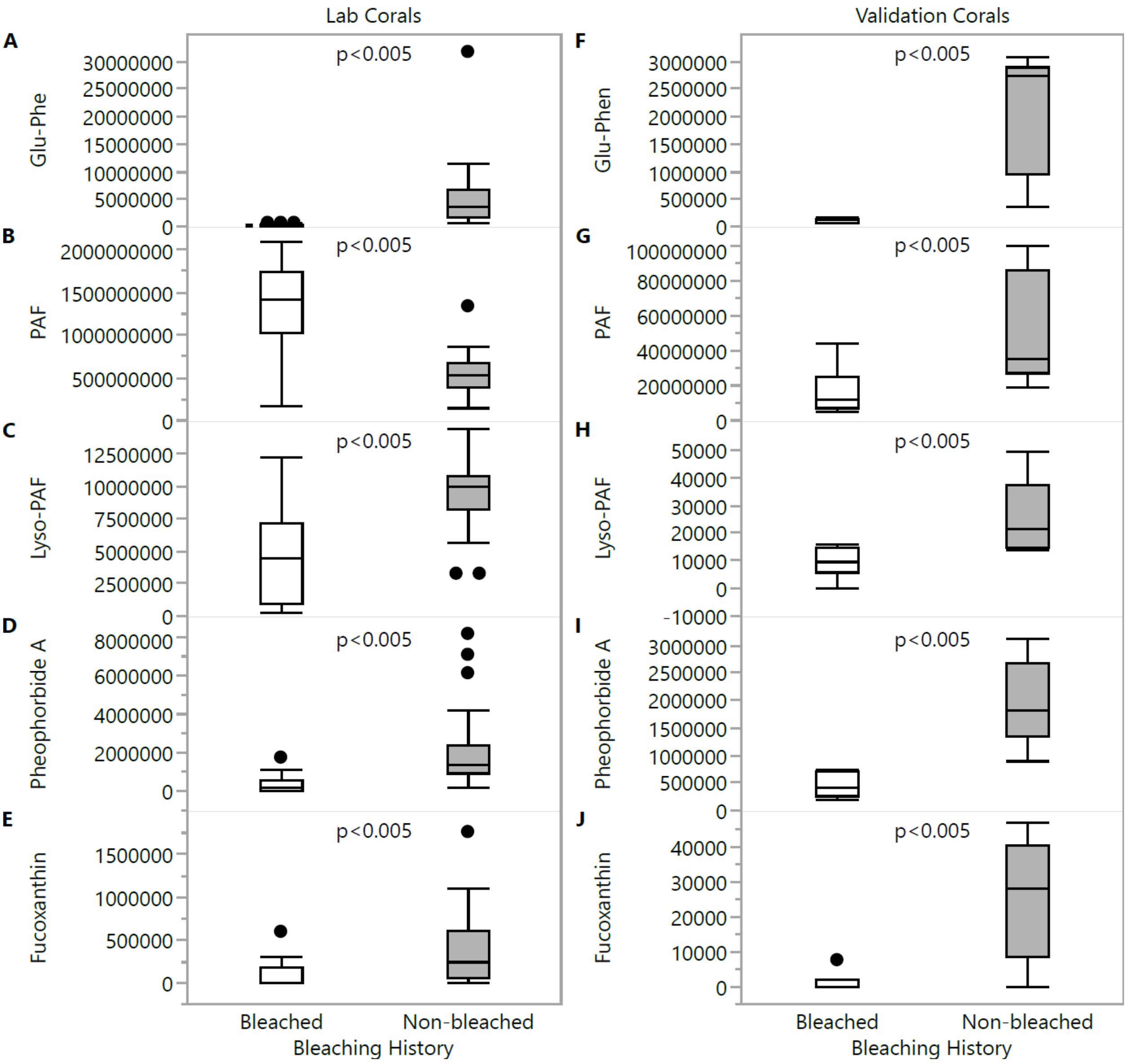


**Supplemental Figure 9.** Box and whisker plots of biologically interesting known metabolites in the ‘lab corals’ (left panels) and the *in situ* ‘validation corals’ (right panels). Boxplots are median with quartiles and whiskers extending 1.5 IQR beyond quartiles.

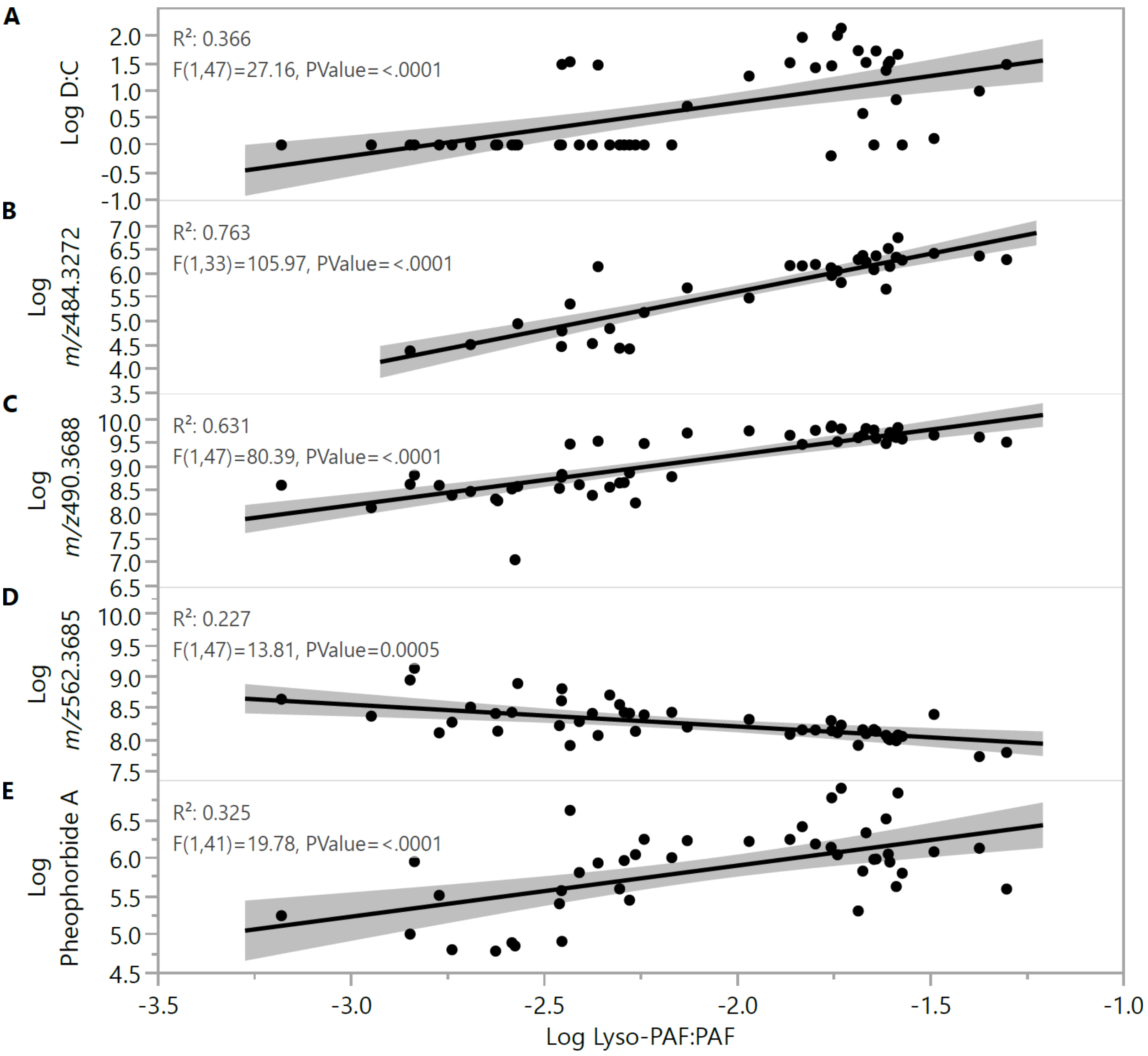


**Figure S10.** Linear regression of the Lyso-PAF:PAF ratio versus symbionts and symbiont-derived metabolites. D:C ratio represents the ratio of *Durisdinum* to *Cladocopium* symbionts. Shaded regions represent the 95% confidence intervals.

**Table S1. Identification and comparison of diacyl betaine lipids in bleached and non-bleached corals.** Measured mass, molecular formulas and PPM error identifying large mass betaine lipids as diacyl forms. The fold difference in abundance of bleached to not-bleached and the p-value from the Wilcoxon Rank-Sum test is also shown. Abbreviations are as follows: B is Bleached, NB is Non-Bleached.

| Measured m/z | Molecular Formula Diacyl | Molecular Formula Monoacyl | PPM error Diacyl m/z | PPM Error Monoacyl | Fold Diff B:NB | p value B vs. NB |
|---|---|---|---|---|---|---|
| ***722.5665*** | $C_{42}H_{76}NO_8$ | $C_{43}H_{80}NO_7$ | 13.02 | 37.34 | 4.1 | <0.00001 |
| ***750.5839*** | $C_{44}H_{80}NO_8$ | NA | 5.99 | >50 | 5.4 | <0.00001 |
| ***774.5832*** | $C_{46}H_{80}NO_8$ | NA | 6.7 | >50 | 7.5 | <0.00001 |
| ***798.5832*** | $C_{48}H_{80}NO_8$ | NA | 6.5 | >50 | 9.9 | <0.00001 |
| ***800.5981*** | $C_{48}H_{82}NO_8$ | NA | 7.42 | >50 | 4.2 | <0.00001 |
| ***822.584*** | $C_{50}H_{80}NO_8$ | $C_{51}H_{84}NO_7$ | 5.34 | 49.29 | 5.5 | <0.00001 |

**Table S2: Family change.** Table of adjusted p values for the molecular families which had Normalized Heat Response significantly different from zero during temperature stress. Each molecular family was tested for each phenotype separately. Abbreviations are as follows: B is Bleached, NB is Non-Bleached, and NS is Not Significant. Bold values indicate significant changes after Benjamani-Hochberg correction.

| Molecular Family | Adjusted p-value (Bleached) | Adjusted p-value (Non-bleached) | Significance |
|---|---|---|---|
| ***Nucleotides*** | **0.003** | 0.866 | Bleached |
| ***Prostaglandins*** | 0.322 | 0.062 | NS |
| ***Peptides*** | 0.279 | 0.286 | NS |
| ***Carnitines*** | **0.006** | **0.037** | Both |
| ***Betaines*** | 0.139 | 0.164 | NS |
| ***Steroids*** | **0.003** | 0.444 | Bleached |
| ***Monoacylglycerides*** | **0.010** | 0.866 | Bleached |
| ***Eicosanoids*** | 0.453 | 0.286 | NS |
| ***Phosphocholines*** | 0.139 | 0.866 | NS |
| ***Fatty Acids*** | **0.001** | 0.866 | Bleached |
| ***Phosphoethanolamines*** | **0.003** | **0.023** | Both |
| ***Phosphatidic Acids*** | 0.215 | 0.722 | NS |
| ***Xanthins*** | **0.027** | 0.866 | Bleached |
| ***Amino Acids*** | 0.443 | 0.866 | NS |
| ***Triterpenoids*** | 0.952 | 0.286 | NS |
| ***Microbial Natural Products*** | 0.152 | 0.740 | NS |
| ***Endocannabinoids*** | 0.757 | 0.286 | NS |
| ***Diterpenoids*** | 0.108 | 0.276 | NS |
| ***Phosphoserines*** | 0.512 | 0.992 | NS |
| ***Indoles*** | 0.829 | 0.701 | NS |
| ***Chlorophyll*** | 0.453 | 0.286 | NS |
| ***Lactones*** | 0.453 | 0.855 | NS |

**Table S3: Family magnitude.** Table of adjusted p values for the magnitude of Normalized Heat Response (NHR) during temperature stress by molecular family. Magnitude of NHR (Euclidean distance from origin) for each family was compared to all other molecules. NS is Not Significant. Bold values indicate significant changes after Benjamani-Hochberg correction.

| Molecular Family | Adjusted p-value | Significance |
|---|---|---|
| ***Nucleotides*** | 0.666 | NS |
| ***Prostaglandins*** | 1.000 | NS |
| ***Peptides*** | 1.000 | NS |
| ***Carnitines*** | 0.189 | NS |
| ***Betaines*** | 0.071 | NS |
| ***Steroids*** | **0.043** | Significant |
| ***Monoacylglycerides*** | **0.000** | Significant |
| ***Eicosanoids*** | 1.000 | NS |
| ***Phosphocholines*** | 1.000 | NS |
| ***Fatty Acids*** | 1.000 | NS |
| ***Phosphoethanolamines*** | 1.000 | NS |
| ***Phosphatidic Acids*** | 0.366 | NS |
| ***Xanthins*** | 0.223 | NS |
| ***Amino Acids*** | 1.000 | NS |
| ***Triterpenoids*** | 1.000 | NS |
| ***Microbial Natural Products*** | 0.471 | NS |
| ***Endocannabinoids*** | **0.043** | Significant |
| ***Diterpenoids*** | 0.223 | NS |
| ***Phosphoserines*** | 1.000 | NS |
| ***Indoles*** | **0.046** | Significant |
| ***Chlorophyll*** | **0.043** | Significant |
| ***Lactones*** | 0.135 | NS |

**Table S4. Outliers.** Table of adjusted p values for the number of outliers in Normalized Heat Response (NHR) to temperature stress by molecular family. Number of outliers in the top 5% of NHR (Euclidean distance from origin) was compared to expectations based on overall proportion. NS is Not Significant. Bold values indicate significant changes after Benjamani-Hochberg correction.

| Molecular Family | Adjusted p-value | Significance |
|---|---|---|
| ***Betaines*** | **0.012** | Significant |
| ***Carnitines*** | 0.367 | NS |
| ***Chlorophyll*** | **0.024** | Significant |
| ***Endocannabinoids*** | 0.588 | NS |
| ***Indoles*** | **0.014** | Significant |
| ***Lactones*** | 0.196 | NS |
| ***Microbial Natural Products*** | 0.532 | NS |
| ***Monoacylglycerides*** | **0.012** | Significant |
| ***Nucleotides*** | 0.367 | NS |
| ***Peptides*** | 0.532 | NS |
| ***Phosphatidic Acids*** | 0.107 | NS |
| ***Phosphocholines*** | 0.107 | NS |
| ***Phosphoethanolamines*** | 0.367 | NS |
| ***Prostaglandins*** | 0.588 | NS |
| ***Steroids*** | **0.024** | Significant |
| ***Xanthins*** | 0.750 | NS |
